# Consolidating the knowledge of the Caatinga’s bryophytes: a comprehensive synthesis with new records

**DOI:** 10.64898/2025.12.19.695594

**Authors:** Jhonyd Jhonata de Oliveira Marmo, Jailton Venilson Ferreira da Silva, Ricely Maria de Moura, Daniel Salgado Pifano, Maria Carolina Tonizza Pereira, Hermeson Cassiano de Oliveira, Mércia Patrícia Pereira Silva

## Abstract

**Introduction:** The Caatinga *sensu lato* (*s.l.*), a Seasonally Dry Tropical Forest, faces significant anthropogenic and climate change pressures. Bryophytes are crucial yet understudied components of this biome, with major knowledge gaps in their distribution, diversity, and conservation status.

**Objectives:** This study aimed to consolidate and expand the knowledge of the Caatinga’s s.l. bryoflora by integrating data from field inventories, herbaria, and literature into a comprehensive occurrence database.

**Methods:** Floristic surveys were conducted in five localities. A dataset was assembled from the three data sources. Multivariate Generalized Linear Models, UPGMA, and Indicator Value analyses compared bryophyte communities among vegetation types. Taxonomic beta diversity was calculated for the entire biome. Conservation analyses assessed species representativeness inside and outside Protected Areas.

**Results:** A total of 627 bryophyte taxa (608 species, 15 varieties, and 4 subspecies), including 54 endemics, was documented for the Caatinga s.l., with five new records added. Ecotone areas showed the highest richness (441 taxa). Bryophyte communities differed among vegetation types, with species turnover as the main driver of beta diversity (0.991). Although 76.4% of all taxa and 64.8% of endemic taxa have at least one record within a Protected Area.

**Conclusions:** The historical perception of low biodiversity in the Caatinga bryoflora is refuted, revealing high diversity across vegetation types. Data gaps in distribution and conservation status are addressed. Strong sampling bias towards Protected Areas likely overestimates in situ protection. Systematic sampling in under-collected regions and stronger conservation policies are urgently needed to safeguard this component of dry forest biodiversity

## Introduction

Biodiversity knowledge forms the foundation for accurately representing natural patterns and processes (Pilowsky *et al*., 2022), thereby supporting effective conservation strategies (Chase *et al*., 2020). However, this understanding is constrained by taxonomic and biogeographical gaps (Linnean and Wallace shortfalls, respectively) (Hortal *et al*., 2015). In light of these gaps, floristic surveys contribute to broadening this understanding and serve as a baseline both for successful conservation strategies (Hanson *et al*., 2023) and for advancing ecological research, particularly in vulnerable ecosystems.

Among the world’s most threatened ecosystems are Seasonally Dry Tropical Forests (SDTFs), as they face intense anthropogenic pressures due to activities such as population growth, grazing, and the demand for energy fuelwood and charcoal (Siyum, 2020). The overlap between these chronic anthropic disturbance and future projections of increased aridity (Allen *et al*., 2017) may drastically reduce the resilience of these forests, affecting their capacity for recovery and the provision of essential ecosystem services (Ribeiro *et al*., 2025). In Brazil, the Caatinga *sensu lato*, a mosaic of SDTF vegetation types within the biome boundaries (hereafter, Caatinga *s.l.*), is projected to experience loss of more than 99% of the woody plant species by 2060, with a trend toward the replacement of narrowly distributed taxa by generalist species (Moura *et al*., 2023). This scenario is exacerbated by estimates that up to 97% of the biome’s territory will lose climatic suitability, where high-altitude areas (800-1000 m) function as essential biodiversity refugia (Teixeira *et al*., 2025).

Regarding bryophytes, floristic inventories in the Caatinga *s.l.* initiated in the 1990s (Pôrto *et al*., 1994; Pôrto *et al*., 1996; Bastos *et al*., 1998a) and substantially advanced floristic knowledge of this biome. More recently, the number of studies is increasing and the bryophytes are a documented component of the Caatinga’s flora (Valente *et al*., 2013; Nascimento *et al*., 2019; Marmo and Silva, 2025a). However, the current knowledge of Caatinga bryophytes remains fragmented. Extensive undersampled areas and significant collection gaps compromise our understanding of their spatial distribution and hinder accurate conservation status assessments (Marmo and Silva, 2025a). Previous compilations, primarily based on herbarium data, have provided preliminary estimates (e.g., Marmo and Silva, 2025a). Despite these advances, no study has yet integrated field inventories, historical literature and herbarium records into a unified, spatially explicit synthesis of bryophyte diversity and conservation in the Caatinga *s.l.* Consequently, their potential response to the impending environmental changes is unassessed. Thus, integrating these scattered records into consolidated databases is crucial for a robust understanding of taxonomic diversity and distribution. According to Cotarelli *et al*. (2025), the inclusion of frequently neglected groups, such as algae, bryophytes, ferns, and lycophytes, alongside angiosperms, is fundamental for biodiversity databases. Such integration ensures a comprehensive ecological perspective of ecosystems, which supports the development and implementation of effective conservation actions.

Therefore, this study aims to provide a modern, comprehensive and up-to-date synthesis of the Caatinga *s.l.* bryoflora by: 1) conducting new floristic inventories in five key areas across the biome; 2) compiling and unifying all available occurrence records from herbaria, literature, and our new collections into a comprehensive database; 3) analysing the floristic composition and beta diversity across major vegetation types; and 4) evaluating the representation of bryophyte species within the Brazilian network of Protected Areas (PAs).

## Materials and methods

### Study area

The Caatinga *s.l.* encompasses the Brazilian northeast Brazil and northern Minas Gerais, totalling an area of 912,529 km². The region is characterised by prolonged droughts, irregular rainfall (400–1200 mm/year), and mean temperatures between 25°C and 30°C (Antunes *et al*., 2022). Vegetation varies according to altitude: in lower and drier areas, it is predominantly deciduous subshrub-shrub vegetation; conversely, higher altitude areas, which receive higher rainfall, support semi-deciduous and evergreen forests, which harbour sub-humid enclaves known as ‘brejos de altitude’ (Moro *et al*., 2024; Alencar *et al*., 2025).

In this study, we followed the vegetation classification system adapted from the Brazilian Institute of Geography and Statistics (IBGE, 2012), replacing the term ‘Savana-Estépica’ (Steppe Savanna) with ‘Caatinga’ — a semi-arid tropical vegetation characterised by shrubs, herbs, thorny woody plants, and Cactaceae — due to contradictions regarding its usage (Bontempo *et al*., 2024) (Figure 1).

**Figure 1.**
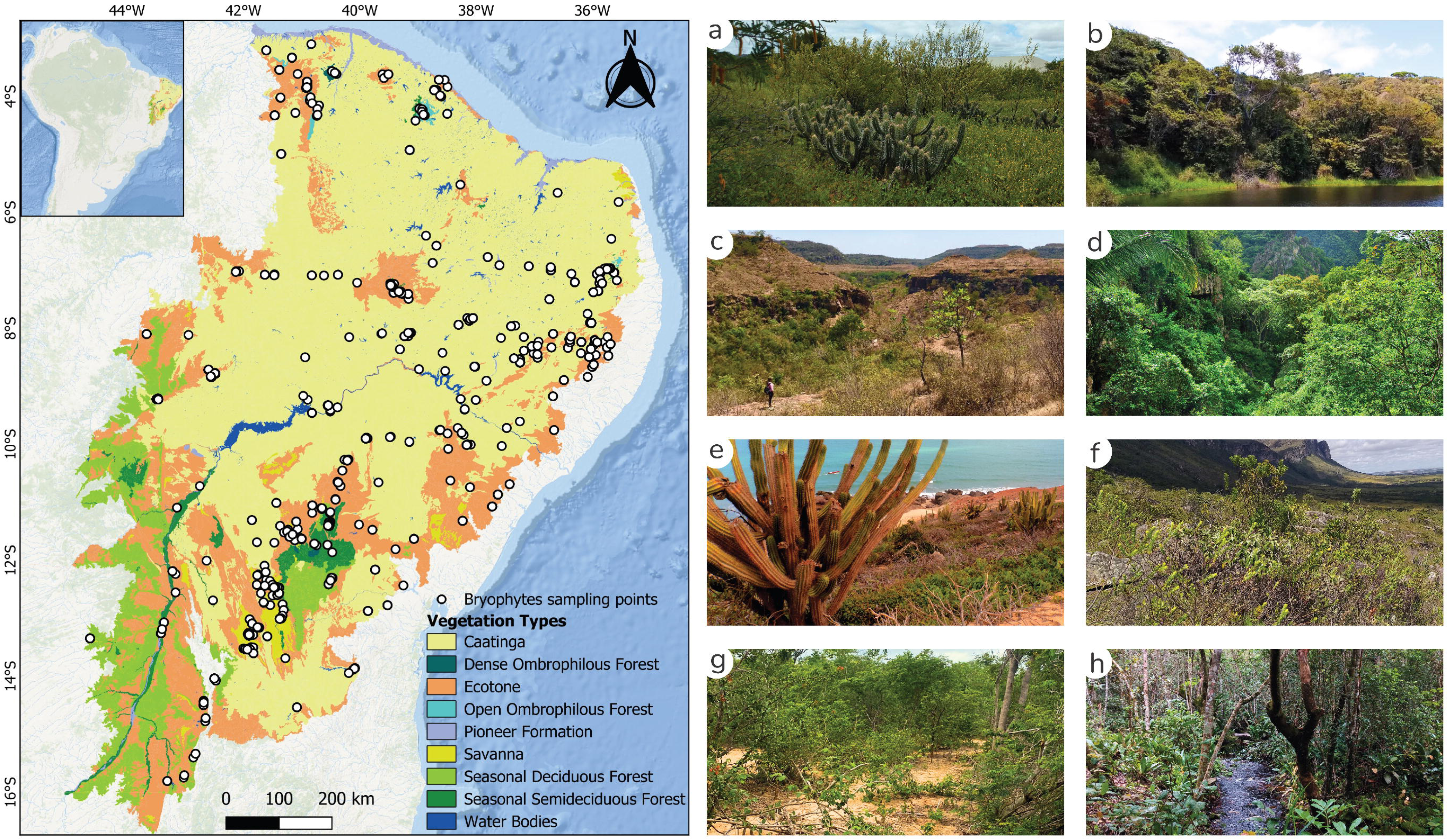
Map of the study area encompassing the Caatinga *sensu lato* and its vegetation types. The map on the left displays the vegetation classification (IBGE, 2012) and the geographic distribution of bryophyte sampling sites (white circles). The right panel (a–h) illustrates the environmental heterogeneity and vegetation types of the region: a) Caatinga; b) Dense Ombrophilous Forest; c) Ecotone (Caatinga-Savanna transition zone); d) Open Ombrophilous Forest; e) Pioneer Formation; f) Savanna; g) Seasonal Deciduous Forest; h) Seasonal Semideciduous Forest. **Source:** Prepared by the author. a) Campus Ciências Agrárias, Petrolina, PE (Photo: Jhonyd Marmo); b) Professor João Vasconcelos Sobrinho Municipal Natural Park, Caruaru, PE (Photo: Alexandre Bittencourt Leite Marques, via Wikimedia Commons, CC BY-SA 4.0); c) Serra de Santo Antônio, Campo Maior, PI (Photo: Maria Elizabeth); d) Ubajara National Park, Ubajara, CE (Photo: Leandro Vargas, via Wikimedia Commons, CC BY-SA 4.0); e) Jericoacoara National Park, Jijoca de Jericoacoara, CE (Photo: Hildebrando de Castro, via Wikimedia Commons, CC BY-SA 4.0); f) Chapada Diamantina National Park, Andaraí, BA (Photo: Tiago Rodrigues Sbarai, via Wikimedia Commons, CC BY-SA 4.0); g) Catimbau National Park, Buíque, PE (Photo: D. Victor Souza); h) Serra da Fumaça, Pindobaçu, BA (Photo: Jhonyd Marmo).

### Floristic survey and specimen processing

Floristic surveys were conducted in five localities selected to represent this vegetation heterogeneity, encompassing distinct floristic compositions (Table 1). Fieldwork was performed during the rainy seasons, between September 2021 and December 2023. Bryophytes were collected through exploratory walks across all accessible substrates (soil, rock, bark, decaying wood, and leaves). Methods for collection, specimen processing, and preservation of botanical samples were adapted from Gradstein *et al*. (2001). All samples were georeferenced using a GPS device (datum SIRGAS 2000). The collected material was deposited in the Herbário de Referência do Sertão Nordestino (HRSN/UNIVASF) and the Herbário Geraldo Mariz (UFP/UFPE). Voucher numbers are listed in Supplemental Material 1.

**Table 1.** Geographic and environmental characteristics of the surveyed areas within the Caatinga *s.l.*, northeastern Brazil.

| Locality | Geographic Coordinates | Vegetation Type (IBGE, 2012) | Climate Type (Köppen) |
| --- | --- | --- | --- |
| Reserva Biológica de Serra Negra | -8.6586, -38.0186 | Seasonal Semideciduous Forest | BSh |
| Serra da Fumaça | -10.6532, -40.3728 | Ecotone | As |
| Sítio Gameleiro | -8.2700, -35.6906 | Open Ombrophilous Forest | As |
| Campus Ciências Agrárias | -9.3311, -40.5490 | Caatinga | BSh |
| Serra de Santana | -10.2754, -40.2138 | Ecotone | As |

**Note:** Vegetation classification follows IBGE (2012); Köppen climate types according to Karger *et al.* (2021).

Taxonomic identification was based on specialized literature. Liverworts (Marchantiophyta) were identified primarily following Gradstein and Costa (2003), and mosses (Bryophyta) using Costa and Pôrto (2022).

### Dataset compilation and spatial processing

To build the occurrence database we integrated three primary sources: 1) new field inventories, 2) herbarium records, and 3) published floristic inventories (see Supplemental Material 1 and 2).

We performed a two-step occurrence validation and distribution assessment through confirmation within the Caatinga biome and assessment of distribution within Brazil and global. First, for compiled records lacking explicit attribution to the Caatinga, occurrence within the target biome was confirmed by consulting the original specimen descriptions in the literature or, when unavailable, by verifying the collection locality via the SpeciesLink platform. The overall distribution pattern of each taxon was assessed for each Brazilian biomes, based on floristic surveys (Carmo and Peralta, 2016; Nascimento *et al*., 2019; Amorim *et al*., 2021; Lima and Peralta, 2021; Remor *et al*., 2021; Aires and Bordin, 2024; Gonçalves *et al*., 2024; Oliveira and Peralta, 2024; Santos *et al.*, 2024) and records from SpeciesLink. Global distribution was checked using the Global Biodiversity Information Facility (GBIF) and Flora e Funga do Brasil (2025), alongside the aforementioned literature sources.

The adopted classification systems were Renzaglia *et al*. (2009) for Anthocerotophyta (hornworts), Goffinet *et al*. (2009) for Bryophyta (mosses) and Crandall-Stotler *et al*. (2009) for Marchantiophyta (liverworts). Nomenclature was updated using The Bryophyte Nomenclator database (Brinda and Atwood, 2025). Taxa recorded as new for the Caatinga *s.l.* were verified against the Flora and Funga do Brasil (2025) database and the SpeciesLink network. Voucher specimens were deposited at the Herbário de Referência do Sertão Nordestino (HRSN/UNIVASF) and the Herbário Geraldo Mariz (UFP/UFPE)

Vegetation types and Protected Areas (PA) vector layers were obtained, respectively, from IBGE (scale 1:250,000) [ibge.gov.br/geociencias/informacoes-ambientais/vegetacao] and the Brazilian National Registry of Conservation Units (CNUC) [https://dados.gov.br/dados/conjuntos-dados/unidadesdeconservacao] (data updated to August 2025). PAs were grouped according to the guidelines of the Brazilian National System of Conservation Units (SNUC) into two management categories: (i) Strict Protection (SP), aimed at nature preservation allowing only the indirect use of natural resources; and (ii) Sustainable Use (SU), which permits environmental use while guaranteeing the sustainability of renewable resources (Lei No. 9.985, 2000).

To mitigate potential cartographic imprecision at the biome boundaries, a 0.03° (∼3.3 km) buffer was applied to the Caatinga *s.l.* polygon [https://terrabrasilis.dpi.inpe.br/downloads/]. Geospatial processing was performed in QGIS (v. 3.44.2). A regular grid with cells of 0.0287° x 0.0287° (∼10 km²) was overlaid on the study area. This resolution was chosen to balance the local aggregation of records with the representation of environmental variability (Marmo and Silva, 2025a). The centroid of each grid cell was used to assign vegetation types and PA categories (Strict Protection or Sustainable Use) to every species occurrence record (see Supplemental Material 3). Grid cells whose centroids did not intersect any PA polygon were classified as ‘Outside Protected Areas’.

### Data analyses

To assess the influence of vegetation type on species composition, we used Multivariate Generalised Linear Models ‘mvabund::manyglm’. This approach is ideal for composition data as it models the mean-variance relationship and handles the dispersion heterogeneity inherent to these datasets (Wang *et al*., 2012). The model was fitted using the binomial family (for presence/absence data) and the ‘cloglog link’ function to address data asymmetry (Wang *et al*., 2025). Model assumptions (log-linearity and mean-variance relationship) were checked visually using Dunn-Smyth residual plots and mean-variance plots (Warton, 2011). Statistical significance was assessed by Analysis of Deviance ‘mvabund::anova.manyglm’, applying Likelihood Ratio Tests with 999 PIT-trap resamples to account for correlation among species (Wang *et al*., 2025). This analysis provided both the global test (assessing significant differences in multivariate composition among groups) and univariate tests, intended to identify which species individually contributed significantly to this general distinction.

Indicator species for each vegetation type were identified using the IndVal method (indicspecies::multipatt), applying the argument ‘func = IndVal.g’ to correct for unequal sizes among sampling groups (Cáceres and Legendre, 2009; Cáceres *et al*., 2025). Floristic similarity was assessed via hierarchical clustering (UPGMA; ‘stats::hclust’), based on a Jaccard dissimilarity matrix ‘vegan::vegdist’, and the dendrogram was constructed using the average linkage method (Oksanen *et al*., 2024). Finally, total beta diversity was calculated using the Jaccard index and partitioned into its turnover and nestedness components ‘betapart::beta.multi’ (Baselga, 2010; Baselga and Orme, 2012).

Taxa were classified based on the incidence of their records within SNUC categories: ‘Inside Protected Areas’ (with at least one occurrence inside a PA) or ‘Outside Protected Areas’ (with all records outside Protected Areas). Among the protected taxa, protection exclusivity was further evaluated, categorising them as: exclusive to SP (records only in this group), exclusive to SU (records only in this group), or shared (with records in both management groups). This assessment was performed for all species and for species endemic to Brazil (Araújo *et al*., 2025). PA coverage richness was summarized for the entire biome.

All statistical analyses were performed in the R (v. 4.5.2) statistical computing environment (R Core Team, 2025). The complete analysis script is available in Supplemental Material 4.

## Results

### Bryophyte diversity of the Caatinga

The floristic surveys recorded a total of 136 taxa (see Supplemental Material 1). Bryophyta was the most representative division with 84 taxa (80 species, 1 subspecies, 3 varieties), distributed in 22 families and 45 genera. The richest families were Leucobryaceae (11), Fissidentaceae (9), and Pottiaceae (9). The division Marchantiophyta accounted for 52 taxa (49 species, 2 subspecies, 1 variety) in 10 families and 26 genera, dominated by Lejeuneaceae (31 taxa), followed by Frullaniaceae (8), and Cephaloziaceae and Ricciaceae (3 taxa each). Five new records for the Caatinga *s.l.* were identified: the species *Calypogeia densifolia* (Steph.) Steph., *Frullania subtilissima* (Nees ex Mont.) Lindenb., *Lejeunea pulchra* C.J. Bastos & Gradst., and *Marchesinia bongardiana* (Lehm. & Lindenb.) Trevis., and the variety *Kurzia capillaris* var. *verrucosa* (Steph.) Pócs (see Supplemental Material 1).

The integration of the new inventories with compiled records resulted in a database of 627 taxa (608 species, 15 varieties, and 4 subspecies), distributed across 76 families and 205 genera (see Supplemental Material 1). Mosses (Bryophyta) were the most representative group (338 taxa), with Fissidentaceae (42 taxa), Leucobryaceae (34), and Sphagnaceae (27) as the richest families. Liverworts (Marchantiophyta, 284 taxa) were dominated by Lejeuneaceae (125 taxa), followed by Lepidoziaceae (22) and Plagiochilaceae (20). Hornworts (Anthocerotophyta) included 5 taxa, represented by the families Notothyladaceae (4 taxa) and Anthocerotaceae (1 taxon) (Figure 2). Regarding endemism, 54 species recorded are endemic to Brazil, three of which are restricted to the Caatinga *s.l.* biome: *Sphagnum contortulum* H.A. Crum, *Sphagnum harleyi* H.A. Crum, and *Riccia bahiensis* Steph.

**Figure 2.**
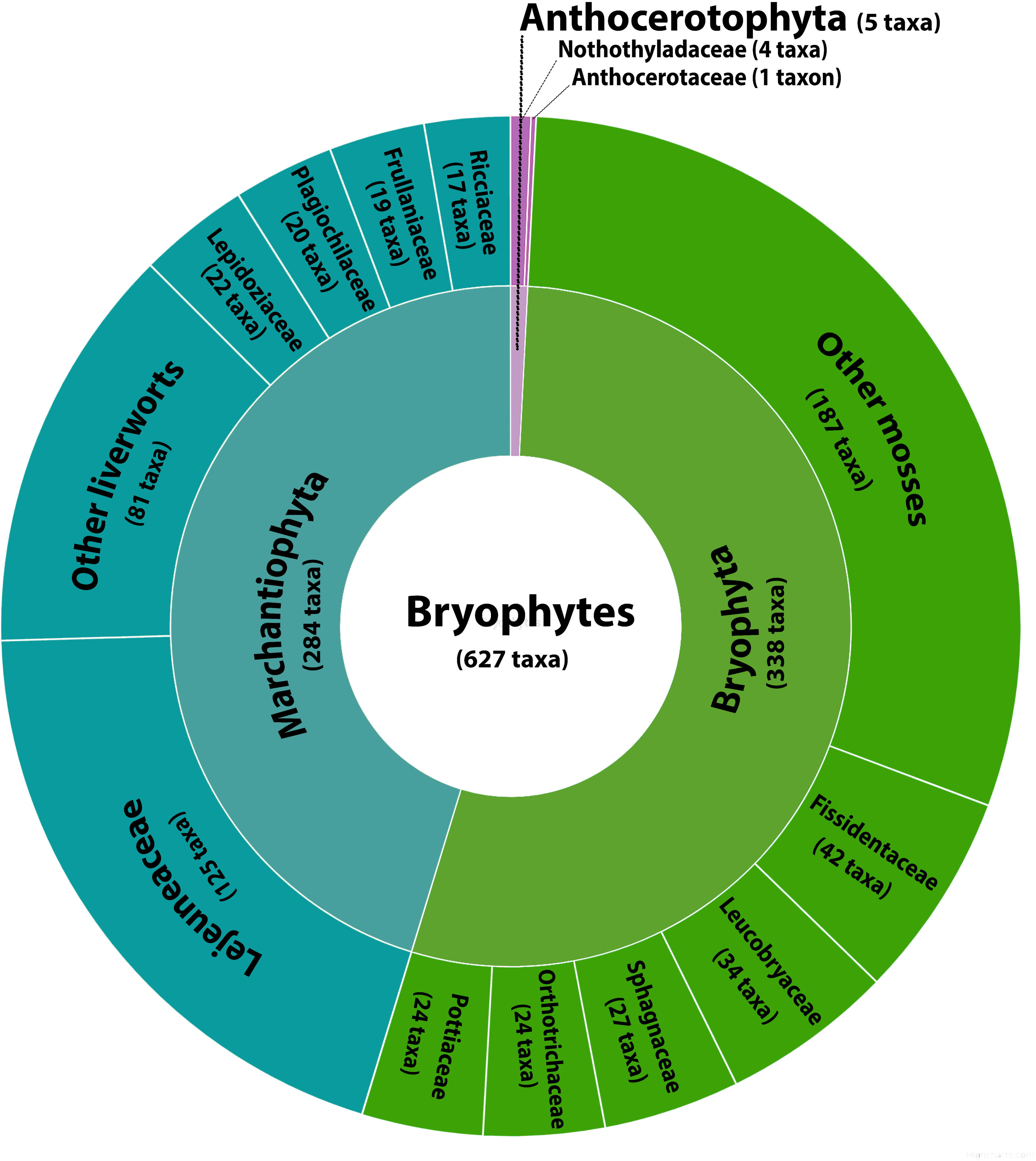
Taxonomic composition of the bryophyte flora of the Caatinga *sensu lato*. The diagram shows the total taxon richness per division (in parentheses) and the relative richness of the most representative families within each group.

Species richness was unevenly distributed across the Caatinga’s *s.l.* vegetation types (Figure 3). Ecotone areas harboured the highest richness (442 taxa, 70.4% of the total). This was followed by Savanna (284), Caatinga (275), Seasonal Semideciduous Forest (207), Open Ombrophilous Forest (145), Seasonal Deciduous Forest (103), and Dense Ombrophilous Forest (63). Pioneer Formations had only a single recorded taxon (Figure 3).

**Figure 3.**
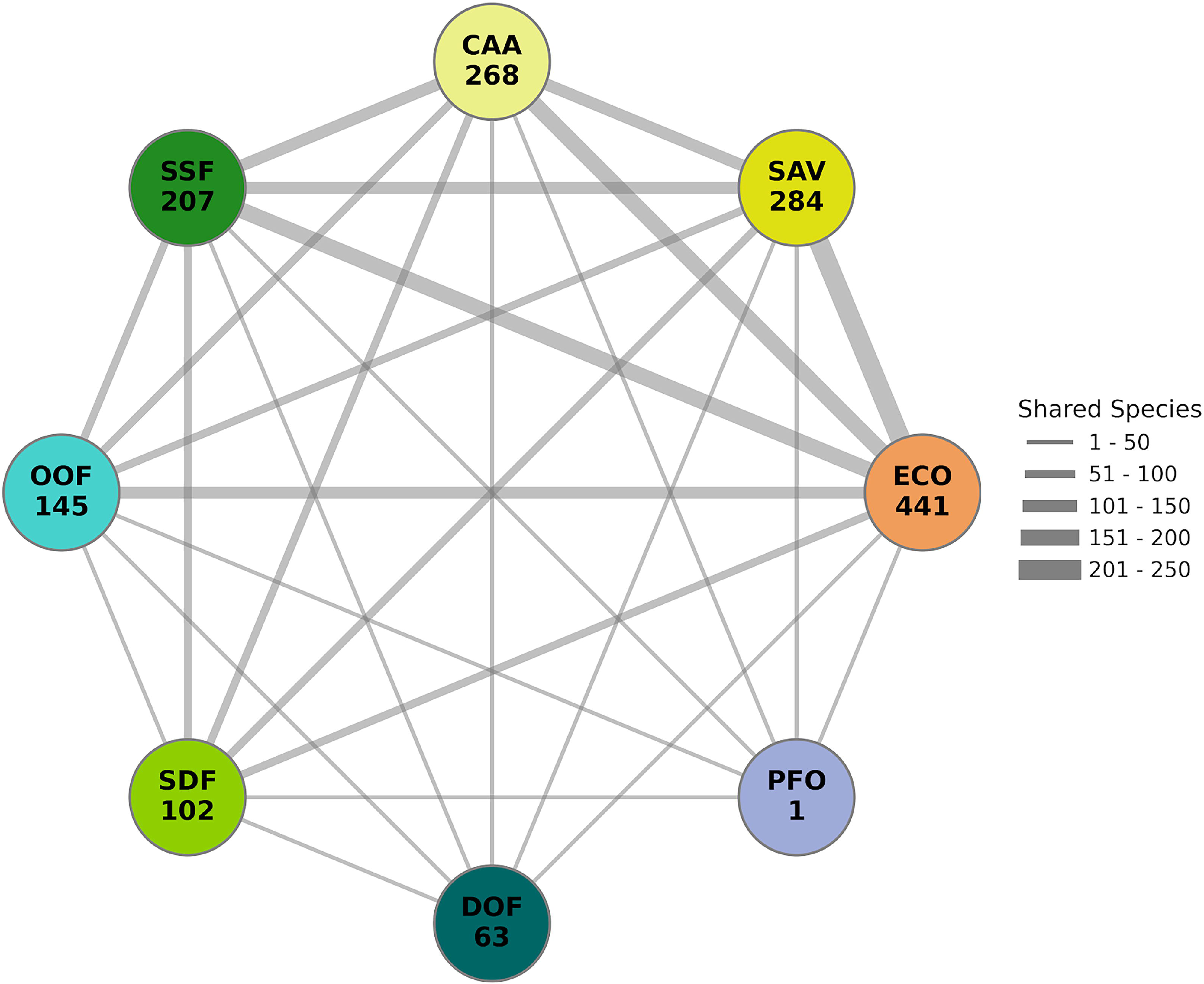
Floristic connectivity network among the main vegetation types of the Caatinga *sensu lato*. Nodes represent vegetation types (see acronyms below) and are sized proportionally to their total species richness. Edges represent the number of shared species between two vegetation types; their thickness is proportional to the strength of the floristic link (see scale). Acronyms: CAA: Caatinga; SDF: Seasonal Deciduous Forest; SSF: Seasonal Semideciduous Forest; ECO: Ecotone; OOF: Open Ombrophilous Forest; DOF: Dense Ombrophilous Forest; PFO: Pioneer Formations; SAV: Savanna.

### Ecological and conservation analyses

#### Ecological analyses

The global ManyGLM confirmed that vegetation type significantly structures bryophyte composition (deviance = 2541, *p-value* = 0.008). Univariate tests identified seven species as key drivers of this compositional differentiation (Table 2). Visual inspection of Dunn-Smyth residuals indicated no patterns of heteroscedasticity among groups. Furthermore, the mean-variance plot confirmed the appropriate fit of the binomial distribution, validating model assumptions for data with low frequency of occurrence (see Supplemental Material 5 and 6).

**Table 2.** Key species driving the floristic differentiation among vegetation types in the Caatinga *s.l*., identified by ManyGLM univariate tests. Species are ordered by their deviance value.

| Species | Deviance | P-value |
| --- | --- | --- |
| *Schlotheimia rugifolia* (Hook.) Schwägr. | 48.65 | 0.001*** |
| *Riccia vitalii* Jovet-Ast | 39.03 | 0.001*** |
| *Syrrhopodon prolifer* Schwägr. | 31.30 | 0.001*** |
| *Frullania brasiliensis* Raddi | 30.15 | 0.002** |
| *Frullania ericoides* (Nees) Mont. | 28.85 | 0.005** |
| *Sphagnum perichaetiale* Hampe | 27.30 | 0.005** |
| *Campylopus arctocarpus* (Hornsch.) Mitt. | 23.39 | 0.023* |

**Note:** \*\*\* $p\text{-value} < 0.001$ ; \*\* $p\text{-value} < 0.01$ ; \* $p\text{-value} < 0.05$ .

O IndVal analysis identified 39 taxa as significant indicators of specific vegetation types, and one indicator species was detected for combinations of two groups (Table 3).

**Table 3.** Significant Indicator Species Analysis (IndVal) for specific vegetation types in the Caatinga *sensu lato*. Only species significantly associated with a single vegetation type are shown (40 taxa total; see Supplemental Material 6 for the complete list). (a) For vegetation types with many indicator species (Dense Ombrophilous Forest and Seasonal Deciduous Forest), only the two species with the highest IndVal values are presented.

| Vegetation Type | Species | IndVal | p-value |
| --- | --- | --- | --- |
| Pioneer Formations | *Riccia vitalii* Jovet-Ast | 0.756 | 0.001*** |
| Dense Ombrophilous Forest (a) | *Octoblepharum pulvinatum* (Dozy & Molk.) Spruce | 0.707 | 0.006** |
|  | *Ceratolejeunea fallax* (Lehm. & Lindenb.) Bonner | 0.500 | 0.009** |
| Savanna | *Leucobryum crispum* Müll. Hal. | 0.551 | 0.043* |
|  | *Pyrrhobryum spiniforme* (Hedw.) Mitt. | 0.476 | 0.048* |
| Seasonal Deciduous Forest (a) | *Tricherpodium beccarii* (Müll. Hal.) Pursell | 0.443 | 0.037* |
|  | *Lejeunea puiggariana* Steph. | 0.316 | 0.037* |
| Open Ombrophilous Forest | *Dibrachiella parviflora* (Nees) X.Q. Shi, R.L. Zhu & Gradst. | 0.331 | 0.048* |
| Seasonal Deciduous Forest + Seasonal Semideciduous Forest | *Zoopsidella integrifolia* (Spruce) R.M. Schust. | 0.327 | 0.049* |

**Note:** \*\*\* *p*-value <0.001; \*\* *p*-value <0.01; \* *p*-value <0.05.

The UPGMA (cophenetic correlation, *r* = 0.972) revealed clear clusters, primarily separating humid forest formation (Dense and Open Ombrophilous Forests) from drier seasonal forests and open vegetation (Figure 4). The partitioning of total beta diversity (Jaccard βtotal = 0.997) showed that species turnover (βsim = 0.991) accounted for almost all dissimilarity, with a negligible contribution from nestedness (βsne = 0.059).

**Figure 4.**
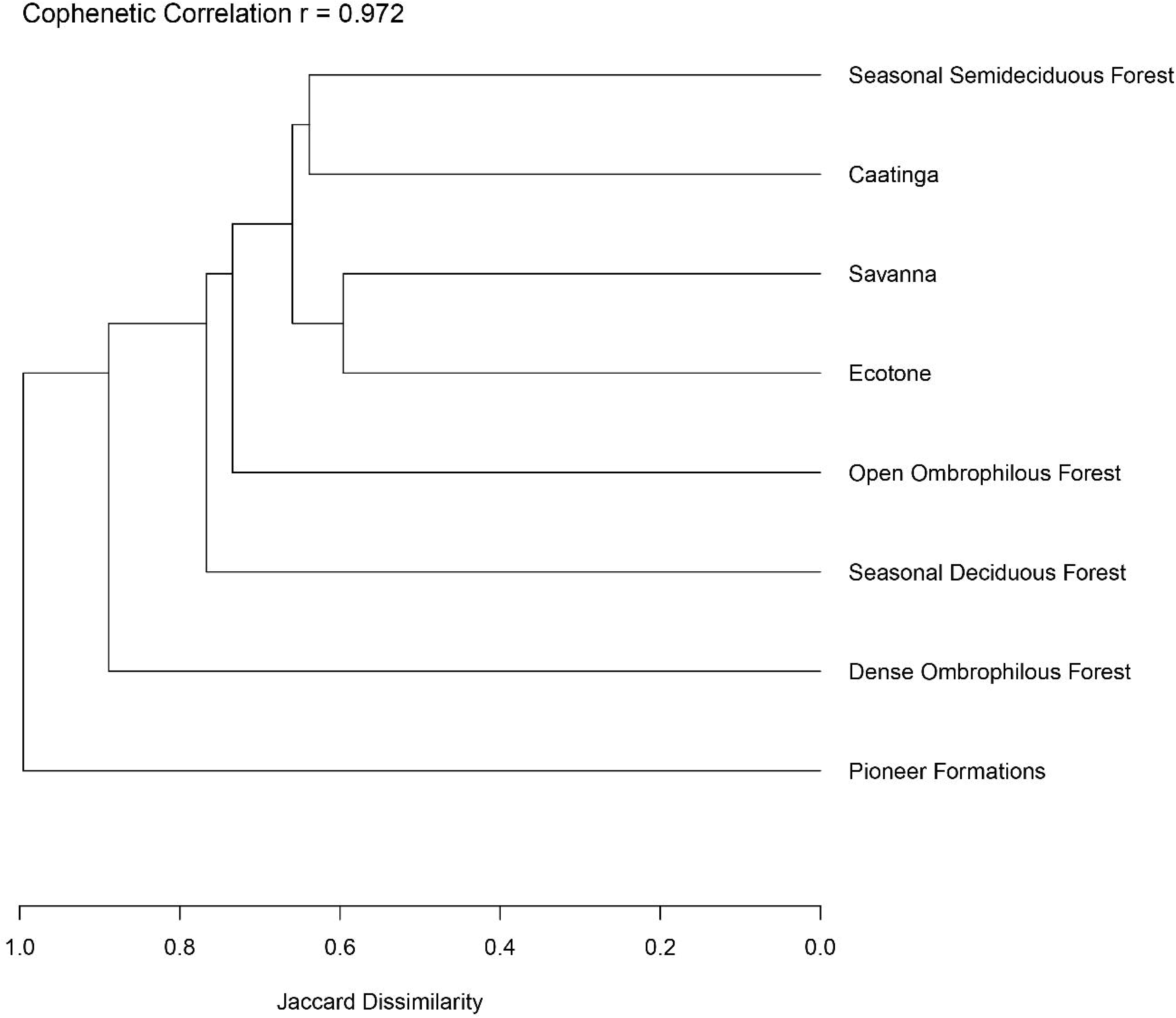
Floristic similarity among vegetation types of the Caatinga *s.l.* based on bryophyte species composition. Dendrogram generated by hierarchical clustering (UPGMA method) using Jaccard dissimilarity.

#### Conservation gap analysis

The spatial distribution of records revealed a pronounced sampling bias, with most collected grid centroids located within or near PAs (Figure 5A). Of the 627 taxa analysed, 479 (76.4%) have at least one occurrence record inside a PA, whereas 148 (23.6%) lack any registered protection (Figure 5B). Among the protected taxa, the largest share (222 taxa, 46.3%) occurs in both Strict Protection (SP) and Sustainable Use (SU) PA categories. Exclusive protection is more common in SP areas (133 taxa, 27.8%) than in SU areas (124 taxa, 25.9%). The PAs with the highest richness were the Parque Estadual das Sete Passagens (179 taxa), Área de Relevante Interesse Ecológico Serra do Orobó (115) and Área de Proteção Ambiental Serra do Barbado (112).

**Figure 5.**
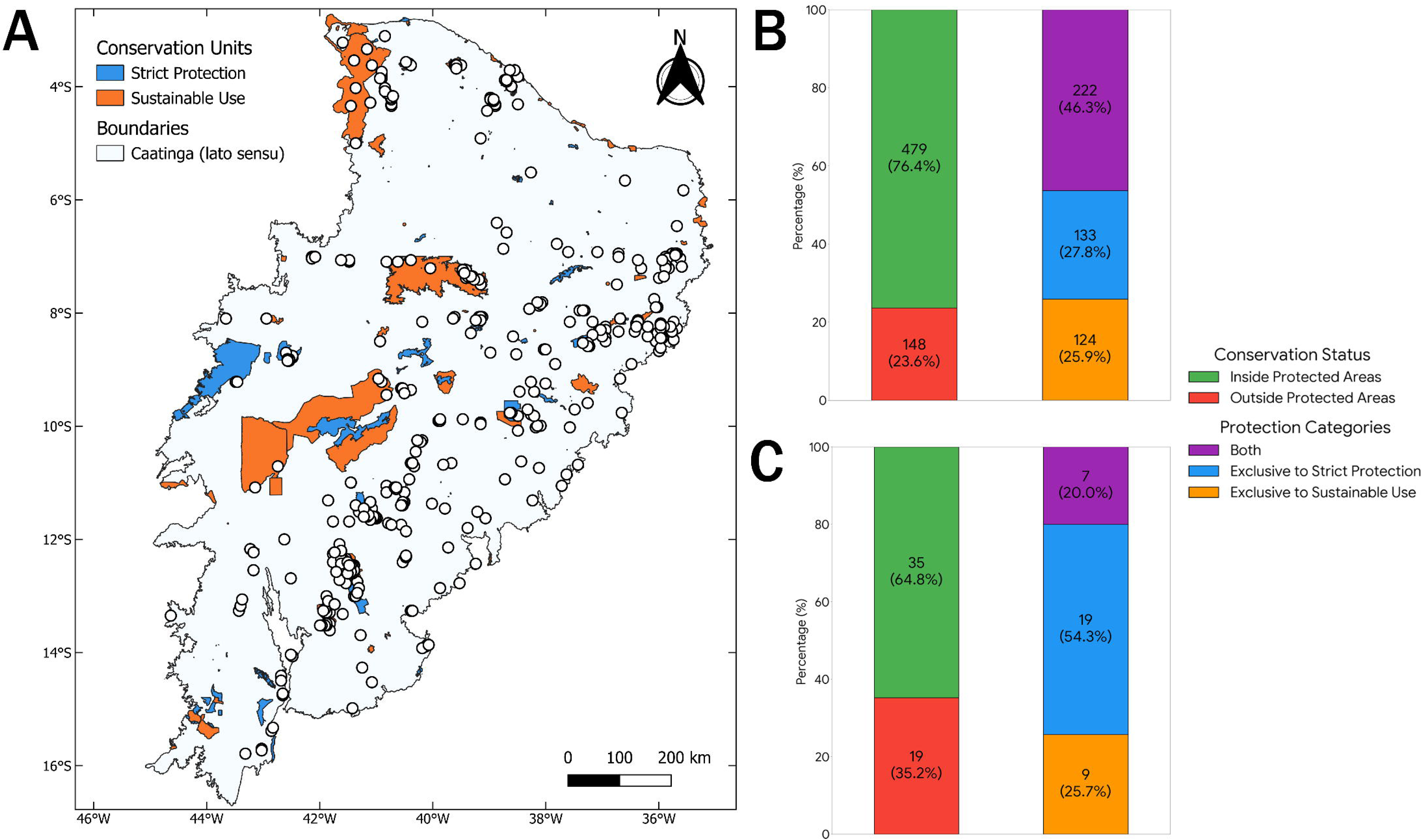
Conservation coverage analysis for bryophytes in the Caatinga *sensu lato*. (A) Spatial distribution of the ∼10 km² grid cells used for analysis, overlaid on the network of Protected Areas (PAs). PA categories: Strict Protection (blue) and Sustainable Use (orange). Red dots indicate grid centroids containing at least one bryophyte record. (B) Proportion of all recorded taxa (n=627) with protection status. ‘Inside Protected Areas’ taxa are subdivided by their occurrence in PA management categories: exclusive to Strict Protection (SP), exclusive to Sustainable Use (SU), or shared between both. (C) Same protection analysis applied only to the subset of taxa endemic to Brazil (n=54).

For the 54 Brazilian-endemic taxa occurring in the Caatinga *s.l.*, the protection scenario is more concerning: 19 species (35.2%) are completely unprotected (Figure 5C). Of the 35 protected endemics, most rely exclusively on SP areas (19 taxa, 54.3%), followed by those in both categories (7 taxa, 20%) and those exclusive to SU areas (9 taxa, 25.7%). The three Caatinga-endemic species show varying status: both *Sphagnum* species are recorded within PAs, while the status of *Riccia bahiensis* (known only from a type specimen collected in 1898) remains unknown.

## Discussion

Our comprehensive synthesis reshapes the understanding of bryophyte diversity in the Caatinga *s.l.* By integrating new fieldwork data with all available records, we documented 627 taxa, a 27.7% increase over the most recent estimate based primarily on herbarium data (Marmo and Silva, 2025a) and surpassing the predicted richness of approximately 608 taxa. This finding refutes the historical perception of the Caatinga as a biome of low bryophyte diversity. Instead, it reveals a rich and complex bryoflora whose distribution is tightly linked to the biome’s environmental heterogeneity.

The concentration of species in Ecotone areas underscores the role of humidity as a driver of bryophyte richness in drylands. These zones, particularly the ‘brejos de altitude’ act as a climatic refugia and as maintainer of regional diversity (Oliveira and Bastos, 2009a, 2009b, 2010a, 2010b; Valente *et al*., 2013; Oliveira and Peralta, 2015). Their high recorded richness is likely a combined result of their suitable environment and sampling bias, as many floristic surveys have been conducted in the humid enclaves. Nevertheless, their high connectivity, evidenced by the network analysis, allows them to share species with both humid forests and drier formations.

Ecologically, bryophytes in the Caatinga *s.l.* are divided into two strategic groups: those from humid areas, whose morphological and reproductive traits are associated with water retention and water-dependent reproductive strategies, such as dioicous reproduction; and those from xeric environments, which exhibit high plasticity for water optimisation and photoprotection (Marmo and Silva, 2025b). Regarding the latter, the presence of traits that aid persistence in these environments suggests a predominance of generalist organisms, as observed in rocky outcrops within the Caatinga *s.l.* (Silva and Germano, 2013). This generalist nature explains the absence of indicator species for the Caatinga vegetation type (Table 3).

The striking predominance of species turnover over nestedness in structuring beta diversity aligns with patterns found in the Caatinga’s woody flora (Apgaua *et al*., 2014) indicating that environmental filtering across different vegetation types selects for distinct species assemblages. Indeed, for bryophytes, environmental filtering exerts a severe influence on distribution, resulting in lower richness in xeric areas due to physiological dependence on water (Geffert *et al*., 2013). Spatially, the structuring of these communities reflects complex phytogeographical patterns and disjunctions shaped by long-distance dispersal capacity (Santos and Costa, 2010), which constrains local occurrence to species possessing morphological and reproductive traits that aid persistence in these environments (Marmo and Silva, 2025b).

This is further supported by the identification of 40 species for specific, mostly humid, vegetation types. The conspicuous absence of indicator species for the Caatinga vegetation type suggests that the bryophytes inhabiting the drier areas are generalists or sun-adapted species with traits for drought tolerance and photoprotection (e.g., costa, papillae, leaf curling; Silva & Germano, 2013; Marmo & Silva, 2025b).

Quantitatively, the Brazilian PA network offers broad coverage for the Caatinga bryoflora, encompassing 76.4% of all recorded taxa and 64.8% of Brazilian-endemic taxa. However, this encouraging statistic is undermined by a pronounced sampling bias, as most historical and recent surveys have been conducted within PAs (Bastos *et al*., 1998b; Nascimento *et al*., 2019; Silva *et al*., 2019; Sousa *et al*., 2024). This bias may inflate protection metrics and mask the true vulnerability of bryophytes in the under-sampled matrix outside PAs, which constitutes over 91% of the biome. Furthermore, when analysing the Caatinga *s.l.* across its entire territory, only 8.9% is under legal protection and of this, a mere 1.8% falls under strict protection categories and approximately 80% corresponds to SU categories (referring to Environmental Protection Areas – APAs, IUCN category V, that permit varying degrees of resource exploitation. This strategy is frequently prioritised by public managers to mitigate land and economic conflicts, as it avoids the need for land expropriation. This vulnerability is aggravated by the sharp disparity in protection coverage among the different states of the region (Teixeira *et al*., 2021). This structural weakness is alarming for habitat specialists and the three Caatinga-endemic species: *Sphagnum contortulum* H.A. Crum, *Sphagnum harleyi* H.A. Crum, and *Riccia bahiensis* Steph. The two *Sphagnum* species are recorded within PAs, whereas the status of *R. bahiensis* remains uncertain, as it is known only from its type material collected in 1898. Despite the existence of PAs near the type locality, the area was recently identified as one of the first aridity nuclei in the region (INPE and Cemaden, 2023), which increases uncertainty regarding the current persistence of the species at this site and it is emblematic of the extinction risks for poorly know taxa.

## Conclusion

In summary, the results obtained in this study redefine the perception of the bryoflora of the Caatinga *s.l.*, refuting the historical paradigm of a bryophyte species-poor biome. By demonstrating the complexity of species composition across the mosaic of vegetation types and presenting a robust database on the distribution and conservation status of the bryoflora, this study fills fundamental knowledge gaps, as discussed by Hortal *et al*. (2015). However, this biome still contains large areas with sampling gaps. Therefore, the intensification of systematic sampling, both within and outside protected areas (PAs), is crucial to correct sampling biases. It is expected that this compiled information will assist in the creation of new PAs, as well as in conservation policies ensuring the protection of the bryoflora, and serve as a basis for future extinction risk assessments for the bryoflora.

## Acknowledgements

We thank the Fundação Coordenação de Aperfeiçoamento de Pessoal de Nível Superior (CAPES) for funding the project through the scholarship granted under process number 88887.806589/2023-00.

## Data availability statement

The data that support the findings of this study are available in the Supplemental Material of this article.

## Supplemental material

**Supplemental Material 1.** Comprehensive database of bryophyte records in the Caatinga sensu lato, integrating data from new field inventories, herbarium collections, and literature, including taxonomic classification, geographical coordinates, and voucher information.

**Supplemental Material 2.** List of data sources and published studies used to compile the comprehensive bryophyte database.

**Supplemental Material 3.** Environmental matrix detailing vegetation types, Protected Areas (PAs), and spatial data associated with the sampling points.

**Supplemental Material 4.** Complete R script used for statistical analyses, including ManyGLM, Indicator Species Analysis (IndVal), and Beta Diversity partitioning.

**Supplemental Material 5.** Summary of statistical outputs, including Global ManyGLM results, Indicator Species Analysis (IndVal), and Beta Diversity partitioning components.

**Supplemental Material 6.** Detailed univariate statistics (deviance and P-values) for each species derived from the ManyGLM model.

## Notes on contributors

**Jhonyd J. O. Marmo:** is a Ph.D. student at the Department of Botany, Federal University of Pernambuco (UFPE), Brazil. His research focuses on the taxonomy, ecology, and distribution of bryophytes. He is particularly interested in the diversity and environmental drivers of the bryoflora in the Brazilian Seasonally Dry Tropical Forests (Caatinga).

**Jailton V. F. Silva:** is a Ph.D. student at the Department of Botany, Federal University of Pernambuco (UFPE), Brazil. His research focuses on the macroecology and diversity patterns of bryophytes. He is particularly interested in the taxonomic, phylogenetic, and functional diversity of bryophytes in Brazil.

**Ricely M. Moura:** is a Master’s student at the Department of Botany, Federal University of Pernambuco (UFPE), Brazil. Her research interests include the taxonomy, ecology, and conservation of bryophytes.

**Daniel S. Pifano:** is a professor at the Federal University of the São Francisco Valley (UNIVASF), Brazil. His research focuses on plant systematics, floristics, and phytosociology. He is particularly interested in forest ecology and the conservation of fragmented landscapes.

**Maria C. T. Pereira:** is a professor at the Federal University of the São Francisco Valley (UNIVASF), Brazil. Her research focuses on ecology and limnology, with a particular interest in the study of aquatic ecosystems.

**Hermeson C. Oliveira:** is a professor at the State University of Piauí (UESPI), Brazil. His research focuses on the taxonomy and ecology of bryophytes. He is particularly interested in taxonomic and ecological investigations of bryophytes from northeastern Brazil, with an emphasis on regional floristic diversity and ecological patterns.

**Mércia P. P. Silva:** is a professor at the Department of Botany, Federal University of Pernambuco (UFPE), Brazil. Her research focuses on the taxonomy, systematics, and ecology of bryophytes. She is particularly interested in the biogeography, spatial distribution, and conservation of the bryoflora in the Brazilian Atlantic Forest.

**Hermeson C. Oliveira:** is a professor at the State University of Piauí (UESPI), Brazil. His research focuses on the taxonomy and ecology of bryophytes. He is particularly interested in taxonomic and ecological investigations of bryophytes from northeastern Brazil, with an emphasis on regional floristic diversity and ecological patterns.

