## Supplemental Material 2 for "Consolidating the knowledge of the Caatinga’s bryophytes: a comprehensive synthesis with new records"

**Floristic Survey List**

Bastos CJP, Albertos B, Bôas SBV. 1998a. Bryophytes from some Caatinga areas in the state of Bahia (Brazil). Tropical Bryology. 14:69–75.

Bastos CJP, Stradmann MTS, Bôas-Bastos SBV. 1998b. Additional contribution to the bryophyte flora of Chapada Diamantina National Park, State of Bahia, Brazil. Tropical Bryology. 15:15–20.

Batista WVSM, Pôrto KC, Santos ND. 2018. Distribution, ecology, and reproduction of bryophytes in a humid enclave in the semiarid region of northeastern Brazil. Acta Botanica Brasilica. 32(2):303–313.

Bôas-Bastos SBV, Bastos CJP, Costa KR. 2017. Brioflora da Área de Relevante Interesse Ecológico Serra do Orobó, municípios de Ruy Barbosa e Itaberaba, Bahia, Brasil. Pesquisas, Botânica. 70:79–98.

Carvalho DN *et al.* 2023. Plants of the Serra do Mucambo: a checklist of a remnant of deciduous seasonal forest within the Caatinga domain in Bahia, Brazil. Brazilian Journal of Botany. 46(4):947–981. <https://doi.org/10.1007/s40415-023-00936-2>.

Correia RP, Nascimento JA, Batista JPS, Silva MPP, Valente EB. 2015. Composição e aspectos de comunidades de briófitas da região da Chapada Diamantina, Brasil. Pesquisas, Botânica. 67:243–254.

Marmo JJO, Silva MPP. 2025a. Bryophytes of a Brazilian seasonally dry tropical forest: an overview of diversity and environmental drivers. *Flora*. 330:152770. <https://doi.org/10.1016/j.flora.2025.152770>

Moraes LA, Araújo MFV, Conceição GM. 2021. Brioflorula (bryophyta\musgos e marchantiophyta\hepáticas) do Parque Estadual Cânion do rio Poti, Buriti dos Montes - PI. Geografia Ensino & Pesquisa. 25:1–42. <https://doi.org/10.5902/2236499447876>.

Nascimento GMG, Conceição GM, Peralta DF, Oliveira HC. 2019. Bryophytes of Serra da Capivara National Park, Piauí, Brazil. Check List. 15(5):833–845. <https://doi.org/10.15560/15.5.833>.

Oliveira HC, Bastos CJP. 2009a. Jungermanniales (Marchantiophyta) da Chapada da Ibiapaba, Ceará, Brasil. Acta Botanica Brasilica. 23(4):1202–1209. <https://doi.org/10.1590/S0102-33062009000400031>.

Oliveira HC, Bastos CJP. 2009b. Antóceros (Anthocerotophyta) e hepáticas talosas (Marchantiophyta) da Chapada da Ibiapaba, Ceará, Brasil. Rodriguésia. 60(3):477–484. <https://doi.org/10.1590/2175-7860200960302>.

Oliveira HC, Bastos CJP. 2010a. Fissidentaceae (Bryophyta) da Chapada da Ibiapaba, Ceará, Brasil. Brazilian Journal of Botany. 33(3):393–405. <https://doi.org/10.1590/S0100-84042010000300003>.

Oliveira HC, Bastos CJP. 2010b. Musgos pleurocárpicos da Chapada da Ibiapaba, Ceará, Brasil. Acta Botanica Brasilica. 24(1):193–204. <https://doi.org/10.1590/S0102-33062010000100019>.

Oliveira HC, Peralta DF. 2015. Adições à brioflora de musgos acrocárpicos (Bryophyta) do Estado do Ceará, Brasil. Pesquisas, Botânica. 67:37–50.

Oliveira HC, Souza AM, Valente EB. 2019. Bryophyte flora of the Apodi Plateau, Ceará, Brazil. Rodriguésia. 70:e00692018. <https://doi.org/10.1590/2175-7860201970072>.

Pôrto KC, Bezerra MFA. 1996. Briófitas de caatinga: 2. Agrestina, Pernambuco, Brasil. Acta Botanica Brasilica. 10(1):93–102. <https://doi.org/10.1590/S0102-33061996000100009>.

Pôrto KC, Silveira MFG, Almeida PS. 1994. Briófitas da caatinga 1: Estação Experimental do IPA, Caruaru - PE. Acta Botanica Brasilica. 8(1):77–85. <https://doi.org/10.1590/S0102-33061994000100009>.

Santos JCV, Oliveira HC, Alves MH. 2021. Estudo das briófitas do Bosque Sagrado da Guarita, Bom Princípio do Piauí, Piauí, Brasil. Research, Society and Development. 10(5):e32710513433. [https://doi.org/10.33448/rsd-v10i5.13433](https://www.google.com/search?q=https://doi.org/10.33448/rsd-v10i5.13433).

Santos ME, Peralta DF, Amélio LA, Fagundes ACA. 2024. Briófitas da Caatinga: conhecendo a biodiversidade de Briófitas da Serra Barra do Vento, Serrinha, Estado da Bahia, Brasil. Hoehnea. 51:e652023. <https://doi.org/10.1590/2236-8906e652023>.

Silva JB, Albuquerque KEA, Trovão DMBM, Lopes SF. 2024. Bryofloristic diversity and conservation value of a protected area in the Brazilian semi-arid region. Phytotaxa. 647(2):159–176. <https://doi.org/10.11646/phytotaxa.647.2.3>.

Silva JB, Germano SR, Maciel-Silva AS, Santos ND. 2019. A small elevational gradient shows negative bottom-to-top bryophyte richness in a seasonally dry forest in Brazil. Cryptogamie, Bryologie. 40(17):219–231. [https://doi.org/10.5252/cryptogamie-bryologie2019v40a17](https://www.google.com/search?q=https://doi.org/10.5252/cryptogamie-bryologie2019v40a17).

Silva JB, Germano SR. 2013. Bryophytes on rocky outcrops in the Caatinga biome: a conservationist perspective. Acta Botanica Brasilica. 27(4):827–835. <https://doi.org/10.1590/S0102-33062013000400023>.

Silva JB, Santos ND, Pôrto KC. 2014. Beta-diversity: effect of geographical distance and environmental gradients on the rocky outcrop bryophytes. Cryptogamie, Bryologie. 35(2):133–163. [https://doi.org/10.7872/cryb.v35.iss2.2014.133](https://www.google.com/search?q=https://doi.org/10.7872/cryb.v35.iss2.2014.133).

Silva TO, Silva MPP, Pôrto KC. 2014. Briófitas de afloramentos rochosos do Estado de Pernambuco, Brasil. Boletim do Museu de Biologia Mello Leitão (N. Sér.). 36:85–100.

Sousa MEB, Valente EB, Oliveira HC. 2024. Brioflora do Parque Nacional Serra das Confusões, Piauí, Brasil. Scientia Plena. 20(2):021201. [https://doi.org/10.14808/sci.plena.2024.021201](https://www.google.com/search?q=https://doi.org/10.14808/sci.plena.2024.021201).

Souza ERF, Silva JB, Pinto AS, Lopes SDF. 2021. An updated checklist of bryophytes for the state of Paraíba, a Brazilian hotspot: new records and biological spectrum in a Seasonally Dry Tropical Forest fragment. Phytotaxa. 516(3):223–236. <https://doi.org/10.11646/phytotaxa.516.3.2>.

Souza MMA. 2023. Brioflora da Serra de Bodopitá (Paraíba) [Bachelor's thesis]. Campina Grande: Universidade Estadual da Paraíba.

Valente EB, Pôrto KC, Bastos CJP. 2013. Species richness and distribution of bryophytes within different phytophysiognomies in the Chapada Diamantina region of Brazil. Acta Botanica Brasilica. 27(2):294–310. [https://doi.org/10.1590/S0102-33062013000200008](https://www.google.com/search?q=https://doi.org/10.1590/S0102-33062013000200008).

**Biomes and mundial distribuition references:**

Aires ET, Bordin J. 2024. Avanços no conhecimento da Brioflora do Pampa brasileiro. Hoehnea. 51:e762023. <https://doi.org/10.1590/2236-8906e762023>

Alonso M, Jiménez JA, Cano MJ. 2019. Taxonomic revision of *Chionoloma* (Pottiaceae, Bryophyta). Annals of the Missouri Botanical Garden. 104(4):563–632. <https://doi.org/10.3417/2019381>.

Amélio LA, Peralta DF, Carmo DM. 2019. Briófitas do Parque Estadual de Campos do Jordão, Estado de São Paulo, Brasil. Hoehnea. 46:e962018. [https://doi.org/10.1590/2236-8906-96/2018](https://www.google.com/search?q=https://doi.org/10.1590/2236-8906-96/2018).

Amélio LA, Peralta DF. 2020. The genus *Notothylas* (Notothyladaceae, Anthocerotophyta) in Brazil. Brazilian Journal of Botany. 43(2):331–340. <https://doi.org/10.1007/s40415-020-00602-x>.

Amorim ET, Menini-Neto L, Luizi-Ponzo AP. 2021. An overview of richness and distribution of mosses in Brazil. Plant Ecology and Evolution. 154(2):183–191. [https://doi.org/10.5091/plecevo.2021.1635](https://www.google.com/url?sa=E&source=gmail&q=https://doi.org/10.5091/plecevo.2021.1635).

Bastos CJP, Silva FVS. 2023. Notas sobre a ocorrência de *Cheilolejeunea savannae* L.P. Macedo, Ilk.-Borg. & C.J. Bastos e *C. intertexta* (Lindenb.) Steph. no Brasil, e restabelecimento de *Cheilolejeunea compacta* (Steph.) M.E. Reiner (Lejeuneaceae, Jungermanniidae). Hoehnea. 50:e542022. <https://doi.org/10.1590/2236-8906e542022>.

Bastos CJP. 2012. Taxonomia e distribuição de *Cheilolejeunea aneogyna* (Spruce) A. Evans (Lejeuneaceae, Marchantiophyta). Acta Botanica Brasilica. 26(3):709–713. <https://doi.org/10.1590/S0102-33062012000300021>.

Bastos CJP. 2017. O gênero *Cheilolejeunea* (Spruce) Steph. (Lejeuneaceae, Marchantiophyta) nas Américas. Pesquisas, Botânica. 70:5–78.

Batista WVSM, Pôrto KC, Santos ND. 2018. Distribution, ecology, and reproduction of bryophytes in a humid enclave in the semiarid region of northeastern Brazil. Acta Botanica Brasilica. 32(2):303–313. <https://doi.org/10.1590/0102-33062017abb0339>.

Bischler-Causse H, Gradstein SR, Jovet-Ast S, Long DG, Salazar Allen N. 2005. Marchantiidae. Flora Neotropica Monograph. 97:1–267.

Bordin J, Yano O. 2013. Fissidentaceae (Bryophyta) do Brasil. Boletim do Instituto de Botânica. 22:1–168.

Canestraro BK, Peralta DF. 2022. Synopsis of *Anomobryum* and *Bryum* (Bryaceae, Bryophyta) in Brazil. Acta Botanica Brasilica. 36:e2021abb0283. <https://doi.org/10.1590/0102-33062021abb0283>.

Carmo DM, Lima JS, Silva MI, Amélio LA, Peralta DF. 2018. Briófitas da Reserva Particular do Patrimônio Natural da Serra do Caraça, Estado de Minas Gerais, Brasil. Hoehnea. 45(3):484–508. <https://doi.org/10.1590/2236-8906-35/2018>.

Carmo DM, Peralta DF. 2016. Survey of bryophytes in Serra da Canastra National Park, Minas Gerais, Brazil. Acta Botanica Brasilica. 30(2):254–265. <https://doi.org/10.1590/0102-33062015abb0235>.

Carvalho DN *et al.* 2023. Plants of the Serra do Mucambo: a checklist of a remnant of deciduous seasonal forest within the Caatinga domain in Bahia, Brazil. Brazilian Journal of Botany. 46(4):947–981. <https://doi.org/10.1007/s40415-023-00936-2>.

Costa DP. 2010a. Briófitas. In: Forzza RC (Ed.), Catálogo de plantas e fungos do Brasil, vol. 1. Rio de Janeiro: Jardim Botânico do Rio de Janeiro, pp. 452–521.

Costa DP. 2016. A synopsis of the family Pottiaceae in Brazil. Phytotaxa. 251(1):1–69. [https://doi.org/10.11646/phytotaxa.251.1.1](https://www.google.com/search?q=https://doi.org/10.11646/phytotaxa.251.1.1).

Dantas FS, Peralta DF. 2025. A new species of the genus *Eccremidium* (Dicranellaceae, Bryophyta) from Brazil. Phytotaxa. 720(2):184–186. [https://doi.org/10.11646/phytotaxa.720.2.10](https://www.google.com/search?q=https://doi.org/10.11646/phytotaxa.720.2.10).

Frahm JP. 1991. Dicranaceae: Campylopodioideae, Paraleucobryoideae. Flora Neotropica Monograph. 54:1–238.

Gonçalves MTA *et al*. 2024. As briófitas da Reserva Particular do Patrimônio Natural Parque das Neblinas, Estado de São Paulo, Brasil: IV Workshop de Briófitas do Brasil. Hoehnea. 51:e512024. <https://doi.org/10.1590/2236-8906e512024>.

Gradstein SR. 1994. Lejeuneaceae: Ptychantheae, Brachiolejeuneae. Flora Neotropica Monograph. 62:1–216.

Gradstein SR. 2015. Annotated key to the species of *Plagiochila* (Marchantiophyta) from Brazil. Pesquisas, Botânica. 67:23–36.

Lima JS, Peralta DF. 2021. Brioflora do Parque Nacional da Serra da Bocaina, de São Paulo, Brasil. Hoehnea. 48:e802020. <https://doi.org/10.1590/2236-8906-80/2020>.

Moura RM. 2025. Atualização taxonômica e guia fotográfico de *Riccia* L. (Marchantiophyta): análise de amostras dos herbários UFP e HRSN [Bachelor's thesis]. Recife: Universidade Federal de Pernambuco.

Nascimento GMG, Conceição GM, Peralta DF, Oliveira HC. 2019. Bryophytes of Serra da Capivara National Park, Piauí, Brazil. Check List. 15(5):833–845. <https://doi.org/10.15560/15.5.833>

Nascimento GMG, Conceição GM, Peralta DF, Oliveira HC. 2019. Bryophytes of Serra da Capivara National Park, Piauí, Brazil. Check List. 15(5):833–845. <https://doi.org/10.15560/15.5.833>.

Oliveira DS, Peralta DF. 2024. Bryophytes from the “Alto da Serra de Paranapiacaba” Biological Reserve, São Paulo - Brazil. Rodriguésia. 75:e01222023. <https://doi.org/10.1590/2175-7860202475073>

Oliveira HC, Bastos CJP. 2009. Jungermanniales (Marchantiophyta) da Chapada da Ibiapaba, Ceará, Brasil. Acta Botanica Brasilica. 23(4):1202–1209. <https://doi.org/10.1590/S0102-33062009000400031>.

Oliveira HC, Souza AM, Valente EB. 2019. Bryophyte flora of the Apodi Plateau, Ceará, Brazil. Rodriguésia. 70:e00692018. <https://doi.org/10.1590/2175-7860201970072>.

Oliveira RR *et al*. 2021. Diversity and conservation of bryophytes in the Amazon-Cerrado transition of Northeastern Brazil. Pesquisas, Botânica. 75:291–313.

Remor D, Pastore JFB, Peralta DF. 2021. Levantamento das briófitas da trilha do Pessegueirinho, Curitibanos, Estado de Santa Catarina, Brasil. Hoehnea. 48:e1182020. <https://doi.org/10.1590/2236-8906-118/2020>.

Santos ME, Peralta DF, Amélio LA, Fagundes ACA. 2024. Briófitas da Caatinga: conhecendo a biodiversidade de Briófitas da Serra Barra do Vento, Serrinha, Estado da Bahia, Brasil. Hoehnea. 51:e652023. <https://doi.org/10.1590/2236-8906e652023>

Silva JVF, Lopes ADT, Oliveira HC. 2024. Briófitas do munícipio de Corrente, Piauí, Brasil. Paubrasilia. 7:e0144. <https://doi.org/10.33447/paubrasilia.2024.e0144>.

Silva JVF, Oliveira HC, Villa PM, Valente EB. 2025. Brioflora epíxila em fragmentos de Floresta Atlântica da Serra da Ibiapaba, Ceará, Brasil. Scientia Plena. 21(6):061201. [https://doi.org/10.14808/sci.plena.2025.061201](https://www.google.com/search?q=https://doi.org/10.14808/sci.plena.2025.061201).
