## Supplemental Material 5 for "Consolidating the knowledge of the Caatinga’s bryophytes: a comprehensive synthesis with new records"

```
# RESULTS

### GLOBAL MANY GLM
Analysis of Deviance Table

Model: mva_dat ~ vegetations_type

Multivariate test:
                 Res.Df Df.diff  Dev Pr(>Dev)
(Intercept)         356
vegetations_type    349       7 2541    0.008 **
---
Signif. codes:  0 ‘***’ 0.001 ‘**’ 0.01 ‘*’ 0.05 ‘.’ 0.1 ‘ ’ 1
Arguments:
Test statistics calculated assuming uncorrelated response (for faster computation)
P-value calculated using 999 iterations via PIT-trap resampling.

## DIAGNOSTICS CHECK
#Residuals Plot (Dunn-Smyth)
```

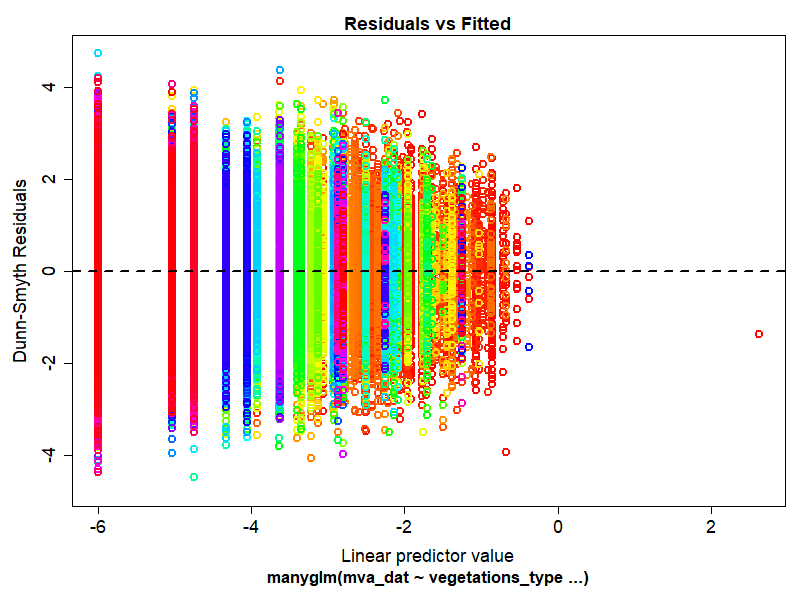

#Mean-Variance Plot

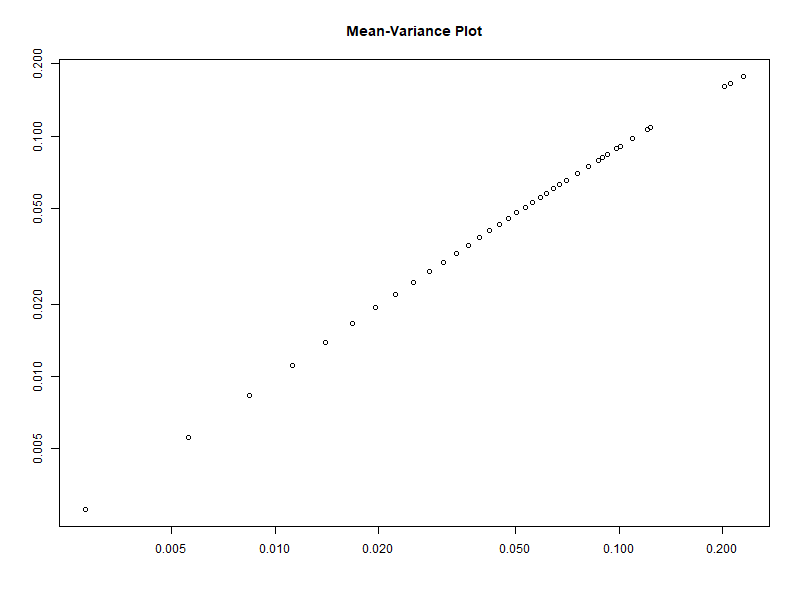

```
### UNIVARIATE MANY GLM

See the Supplementary Data X


### INDVAL

Multilevel pattern analysis
 ---------------------------

 Association function: IndVal.g
 Significance level (alpha): 0.05

 Total number of species: 627
 Selected number of species: 40
 Number of species associated to 1 group: 39
 Number of species associated to 2 groups: 1
 Number of species associated to 3 groups: 0
 Number of species associated to 4 groups: 0
 Number of species associated to 5 groups: 0
 Number of species associated to 6 groups: 0
 Number of species associated to 7 groups: 0

 List of species associated to each combination:

 Group Dense Ombrophilous Forest  #sps.  28
                              stat p.value
Octoblepharum pulvinatum     0.707   0.006 **
Ceratolejeunea fallax        0.500   0.009 **
Prionolejeunea scaberula     0.500   0.009 **
Pteropsiella frondiformis    0.500   0.009 **
Trichosteleum lonchophyllum  0.500   0.009 **
Bryopteris filicina          0.500   0.017 *
Cololejeunea paucifolia      0.500   0.017 *
Lejeunea serpillifolioides   0.500   0.017 *
Orthostichidium pentastichum 0.500   0.017 *
Porotrichum longirostre      0.500   0.017 *
Schwetschkea fabronioides    0.500   0.017 *
Entodon beyrichii            0.494   0.017 *
Metzgeria uncigera           0.494   0.015 *
Leucoloma cruegerianum       0.492   0.017 *
Chryso-hypnum elegantulum    0.492   0.022 *
Macromitrium regnellii       0.492   0.020 *
Crossomitrium patrisiae      0.492   0.027 *
Diplasiolejeunea brunnea     0.492   0.027 *
Lepidopilum muelleri         0.492   0.027 *
Meteorium deppei             0.492   0.027 *
Ceratolejeunea cornuta       0.485   0.018 *
Metzgeria lechleri           0.472   0.024 *
Orthostichopsis tetragona    0.455   0.040 *
Metzgeria ciliata            0.451   0.049 *
Metzgeria bahiensis          0.450   0.029 *
Telaranea diacantha          0.436   0.037 *
Riccardia digitiloba         0.408   0.036 *
Micropterygium lechleri      0.404   0.050 *

 Group Open Ombrophilous Forest  #sps.  1
                         stat p.value
Dibrachiella parviflora 0.331   0.048 *

 Group Pioneer Formations  #sps.  1
                stat p.value
Riccia vitalii 0.756   0.001 ***

 Group Savanna  #sps.  2
                        stat p.value
Leucobryum crispum     0.551   0.043 *
Pyrrhobryum spiniforme 0.476   0.048 *

 Group Seasonal Deciduous Forest  #sps.  7
                                      stat p.value
Tricherpodium beccarii               0.443   0.037 *
Lejeunea puiggariana                 0.316   0.037 *
Fissidens pellucidus var. papillifer 0.316   0.048 *
Micromitrium tenerum                 0.316   0.040 *
Zanderia octoblepharis               0.316   0.042 *
Jonesiobryum termitarum              0.316   0.043 *
Cheilolejeunea savannae              0.316   0.039 *

 Group Seasonal Deciduous Forest+Seasonal Semideciduous Forest  #sps.  1
                          stat p.value
Zoopsidella integrifolia 0.327   0.049 *
---
Signif. codes:  0 ‘***’ 0.001 ‘**’ 0.01 ‘*’ 0.05 ‘.’ 0.1 ‘ ’ 1

### CORRELATION COPHENETIC

[1] 0.972


### BETA TAXONOMIC DIVERSITY

$beta.JTU
[1] 0.9916616

$beta.JNE
[1] 0.005901234

$beta.JAC
[1] 0.9975628
```
